# Defining the heterogeneous molecular landscape of lung cancer cell responses to epigenetic inhibition

**DOI:** 10.1101/2024.05.23.592075

**Authors:** Chuwei Lin, Catherine M. Sniezek, Christopher D. McGann, Rashmi Karki, Ross M. Giglio, Benjamin A. Garcia, José L. McFaline-Figeroa, Devin K. Schweppe

## Abstract

Epigenetic inhibitors exhibit powerful antiproliferative and anticancer activities. However, cellular responses to small-molecule epigenetic inhibition are heterogeneous and dependent on factors such as the genetic background and metabolic state of cells, as well as on-/off-target engagement of individual small-molecule compounds. The molecular study of the extent of this heterogeneity often measures changes in a single cell line. To more comprehensively profile the effects of small-molecule perturbations and their influence on heterogeneous cellular responses, we present a molecular resource based on the quantification of chromatin, proteome, and transcriptome remodeling due to histone deacetylase inhibitors (HDACi) in non-isogenic cell lines. Through quantitative molecular profiling of 10,621 proteins, these data reveal coordinated molecular remodeling of HDACi treated cancer cells. HDACi-regulated proteins differ greatly across cell lines with consistent (JUN, MAP2K3, CDKN1A) and divergent (CCND3, ASF1B, BRD7) cell-state effectors. Together these data provide valuable insight into cell-type driven and heterogeneous responses that must be taken into consideration when monitoring molecular perturbations in culture models. We have also built a web interface for the extensive amount of data to allow users to explore the data as a resource for understanding chemical perturbation of diverse cell types.

## Introduction

Deconvoluting the myriad effects downstream of small molecule perturbations is essential to therapeutic development. Whole proteome and transcriptome analyses of these perturbations can validate and identify proposed mechanisms of action and identify cellular effects for therapeutics and tool compounds. To date, however, many large-scale proteome analyses of drug perturbations have been limited to a single cell line treated with a cohort of small-molecule tool compounds and therapeutic drugs (Supplementary table 1). The selection of cell lines in these studies has been in part a practical consideration as each new cell line included leads to a rapid expansion in the number of samples that must be processed and compared. Unfortunately, this often means sacrificing understanding of the diverse molecular contexts of different cellular models to enable the analysis of larger cohorts of small-molecule compounds (P.-H. Chen et al., 2019).

Large-scale viability screening of small molecule perturbations in the PRISM dataset has shown that drugs targeting the same protein or pathway can drive heterogeneous responses in cancer cells (Corsello et al., 2020). Recent transcriptome efforts for bulk and single-cell samples have also uncovered divergent cellular responses when cells are treated with therapeutic and tool compounds (Srivatsan et al., 2020). At the single-cell level, these studies revealed inconsistent molecular responses of individual, isogenic cells to small molecule perturbation suggesting that additional context and molecular understanding are necessary to dissect the molecular consequences of chemical perturbation. More challenging still is the fact that transcriptomic profiling of cellular perturbations cannot measure the direct protein targets of many of these perturbations (Maltz & Wollman, 2022). Thus, owing to the limited correlation between RNA and protein measurements (Wang et al., 2019), there remains a gap in our knowledge of the proteomic consequences of drug perturbations across multiple, representative cell lines.

Recently, several studies have aimed to bridge this gap. These include the DeepCoverMOA and decryptE datasets (Mitchell et al., 2023; Zecha et al., 2023), which represent some of the largest, publicly-available chemical perturbation-based proteomic datasets. DeepCoverMOA analyzed HCT116 cells in response to 875 therapeutic and tool compounds and revealed that a majority of the measurable proteome was accessible to regulation by small molecule treatment. The resulting proteome remodeling responses were then used to assemble proteomic networks to define consistent chemical mechanisms of actions. In a related study, the decryptE dataset studied dose-dependent chemical responses of 144 compounds in Jurkat cells to determine similarities and differences in cellular mechanisms of action. Yet, an important caveat to these two studies was that the whole proteome analysis to decipher compound mechanisms of action were each done in single, isogenic cell lines which may not be representative of the genetic and molecular heterogeneity when considering non-isogenic cell models (Begley & Ellis, 2012; Raghavan, 2022). Interestingly, both the DeepCoverMOA and decryptE datasets evidenced high overall compound activity for histone deacetylase (HDAC) inhibitors (HDACi) such as vorinostat, quisinostat, and nexturastat (Mitchell et al., 2023; Zecha et al., 2023).

HDAC regulation is key to the cellular chromatin state and is frequently dysregulated in cancer (Y. Li & Seto, 2016) and known to exhibit lineage-specific dependencies in cancer cells (Y. Zhang et al., 2023). The relationship between HDACs and cancer progression has driven a decades-long effort to develop targeted chemical HDAC inhibitors (HDACi) (Yoshida et al., 1990) and FDA approvals for several of these inhibitors to treat lymphomas and melanomas (B. S. Mann et al., 2007; McDermott & Jimeno, 2014; Richardson et al., 2015). While HDAC inhibitors have shown promising preclinical results for many cancers, clinical trial results vary (Mamdani & Jalal, 2020). Vorinostat monotherapy in non-small cell lung cancer (NSCLC) patients did not show objective antitumor activity, but toxicity with adverse effects such as fatigue (Traynor et al., 2009, p. 20). Other phase II monotherapy of romidepsin and pivaloyloxymethyl butyrate resulted in little efficacy in patients, and had side effects such as fatigue, nausea, and dysgeusia (Reid et al., 2004; Schrump et al., 2008).

While comprehensive understanding of the pleiotropic molecular consequences remains elusive (Miyanaga et al., 2008), treatment of cells with HDACi induces expression of p21/CDKN1a leading to cell cycle arrest (Richon et al., 1996; Xiao et al., 1999), alters expression of c-Jun (Vrana et al., 1999) and the apoptotic regulators Bcl-2 and Bcl-xL (X. X. Cao et al., 2001; Fandy & Srivastava, 2006), attenuates AKT/mTOR signaling leading to autophagy (Y.-L. Liu et al., 2010), and suppresses IFN-mediated signaling (Shulak et al., 2014). Recent chemoproteomics analyses have also revealed that heterogeneous responses to HDAC inhibition may be driven in part by the abundance of HDAC protein complex members and the engagement of off-target HDACi binders such as MBLAC2 (Lechner et al., 2022). Thus, these effects have been shown to be both cell-type dependent and cell-type independent, further confounding our understanding of the molecular consequences of HDACi cellular treatments.

Here, we set out to determine the effects of the HDACi treatment on proteome remodeling in non-isogenic cell lines. In particular we focused on cell treatments with well-established HDACi compounds to build on previous datasets such as DeepCoverMOA and decryptE—including, belinostat, CUDC-101, trichostatin A (TSA), panobinostat, abexinostat and vorinostat (SAHA). We did this in the context of a genetically diverse panel of lung cancer cell lines with KRAS, EGFR, TP53, CDKN1A, STK11 mutations and integrated proteomic, phosphoproteomic, and transcriptomics to explore drivers of the heterogeneous HDACi responses in these cells. Quantification of histone modifications status in multiple cell lines and thermal stability analyses then allowed us to determine the molecular linkage and effects of on-target and off-target protein engagement with HDAC inhibitors. We have made these data available as a resource (https://github.com/SchweppeLab/HDAC-Perturbation-DataViewer) to understand how chemical perturbation analyses are affected by preclinical cell line choice and exploration of how cellular diversity affects the characterization of small molecule compound mechanisms of action.

## Results

### Cell line selection for HDACi perturbation analyses

We selected five cell lines for the analysis of cell-type specific proteome profiling. Cell line selection was based on three factors: presence of comparative datasets for benchmarking of the resulting quantitative proteomics analyses, representative mutational status of non-small cell lung cancers (NSCLCs), and divergent responses to epigenetic perturbation in the PRISM drug repurposing screen (Tsherniak et al., 2017). Based on these criteria, we chose HCT116, A549, PC9, H292, and PSC1 cells as representative cancer cell models to measure the effects of HDACi perturbations.

Two KRAS mutant lines were included—HCT116 (KRAS G13D colon cancer cells) and A549 (KRAS G12S lung adenocarcinoma)—as they have previously been characterized in proteomic and transcriptomic perturbation analyses (Supplementary table 1)(Mitchell et al., 2023; Nusinow et al., 2020; Srivatsan et al., 2020; Tsherniak et al., 2017). For the remaining cell models we focused on NSCLC cell lines as this cancer subtype presents divergent preclinical and clinical responses to HDACi treatment (Mamdani & Jalal, 2020). The three additional lung cancer cell lines were chosen for their representative mutational statuses: H292 is a mucoepidermoid carcinoma line (CRTC1-MAML2 fusion, NF2 P496Tfs), PC9 is a adenocarcinoma line (EGFR E746_A750del, TP53 R248Q), and PSC1 is a doxorubicin-resistant derivative of the INER-51 adenocarcinoma cell line (P-gp/MDR-1 negative; Supplementary table 2).

We verified that our selected panel of cell lines was representative of distinct phenotypic responses to small-molecule perturbation. To assess this, we used the PRISM drug repurposing screen, which provides a comprehensive reference of cell viability responses across 911 diverse cell models (Corsello et al., 2020). We calculated the mean viability fold change for all 911 model cell lines in PRISM. We then compared the global mean of cell viability in response to each drug to the viability for the individual cell lines in our selected panel (A549, PC9, A549, HCT116) as well as the composite mean viability of all four overlapping cell lines. The composite mean of our four cell lines better recapitulated the global PRISM data viability data (four cell lines: r_Pearson_ = 0.836) than any one of our cell lines (PC9: r_Pearson_ = 0.706; HCT116: r_Pearson_ = 0.702; H292: r_Pearson_ = 0.625; A549: r_Pearson_ = 0.593, Supplementary Figure 1A).

### Focusing on HDAC inhibition in cancer cell lines

Proteomic studies of the responses of cells to small-molecule drug perturbations have been a powerful means to understand diverse mechanisms of action for therapeutic and tool compounds (Eckert et al., 2024; Mitchell et al., 2023; Zecha et al., 2023). To understand how cancer cell lines with non-isogenic backgrounds respond to pleiotropic effects of drug perturbations we sought to measure proteomic remodeling of multiple cancer cell lines with diverse, and representative, genetic backgrounds in the context of lung cancer—including KRAS, EGFR, TP53, CDKN1A, and STK11 mutations (Figure 1A). We focused on HDAC inhibition (Eckert et al., 2024; Zecha et al., 2023), as epigenetic inhibition is a promising target across multiple malignancies and based on the strength of proteome remodeling previously reported in proteomic (decryptE and DeepCoverMOA) and transcriptomic (Sci-plex) datasets (Mitchell et al., 2023; Srivatsan et al., 2020; Zecha et al., 2023).

We selected six HDAC inhibitors with known canonical and off-target effects, including belinostat (pan-HDACi), CUDC-101 (pan-HDACi, HER2/EGFR inhibitor) (Y. Li & Seto, 2016; Shanmugam et al., 2022), trichostatin A (TSA, pan-HDACi), panobinostat, abexinostat (HDAC1/2/3/6/8/10 inhibitor) (Buggy et al., 2006), and vorinostat (pan-HDACi). To ensure that we could robustly capture proteome remodeling effects at the cell-type- and compound-specific level, we also treated these cell lines with the p38 kinase inhibitor ralimetinib as a non-HDACi-treated outgroup. We performed viability assays and dose response curves to measure the cellular effects of HDAC inhibition (Figure 1B, Supplementary Figure 1B). We found that a drug concentration of 10 μM generally maintained a viability ratio compared to DMSO treatment controls greater than 50% viability after treatment. This was consistent with clinical observation of HDAC inhibitors in patient plasma at levels up to 8±4 μM (Kelly et al., 2005).

Owing to the clinical concentrations and collection of previous benchmarking perturbation proteomics datasets, we performed our proteomics analyses at 10 μM of HDACi for 24 hr. At this time point histone acetylation increased with HDAC treatment in H292 and A549 cells and the relative abundance of histone methylation events decreased (Figure 1C). As expected, each of the cell lines was differentially sensitive toward the seven tested compounds (Supplementary Figure 1B). PC9 cells were highly sensitive to HDACi and ralimetinib treatment, especially CUDC-101 (Figure 1B). The sensitivity to CUDC-101 was likely driven by the known off-target inhibition of EGFR by CUCD-101 as PC9 cells harbor a deletion of E746-A750 in EGFR and EGFR gene amplification (Nukaga et al., 2017). Additionally, A549 cells were sensitive to belinostat to a greater extent than the other HDAC inhibitors. Conversely, PSC1 cells were largely insensitive to HDACi and ralimetinib treatment.

### Proteome quantification of cancer cell responses to HDAC inhibition

Proteome remodeling due to HDACi was measured in cancer cells treated with 10 µM of each compound or vehicle control (DMSO) for 24 hours prior to proteomic and viability assays. In total, we performed quantitative proteomic analysis of the 35 single-perturbation, cell-by-drug combinatorial treatments (Figure 1A). We quantified the relative abundance of proteins in comparison of vehicle (DMSO) controls for each of these perturbations and replicate analyses generated highly correlated proteome abundance shifts (Figure 1A). In total, we acquired quantitative measurements for 476,387 peptides, 6,611 phosphorylation sites, 177 histone sites, and 10,237 unique proteins. Protein quantification between biological replicates was highly reproducible with a median Pearson correlation (r) for biological replicates of each drug and cell line of 0.996 (Supplementary Figure 1C). The median coefficient of variation for protein measurements between biological replicates was 4.4% (Supplementary Figure 1D).

Of 10,237 total proteins we quantified, 6,145 proteins were found in all five cell lines (Supplementary Figure 1E), 8,083 proteins were found in at least three cell lines, and 1,311 were found in only one cell line (Supplementary Figure 1F). The six HDACi and ralimetinib all have the potential to drive pleiotropic remodeling of proteins and protein complexes (Mader et al., 2008; Mamdani & Jalal, 2020). Interestingly, though the HDACi treatment drove strong proteome remodeling, PCA analysis clustered cell-line-by-drug groups based predominantly on cell lines, not the compound used to treat these cells. The one notable difference was that compared to matched vehicle controls in each cell line, ralimetinib-treated cells clustered together in principal component space irrespective of the cell line (Supplementary Figure 1G).

### Comparison of cell-type specific proteome responses to benchmarking datasets

Comparison of our proteomic data to the DeepCoverMOA dataset in HCT116 cells (Mitchell et al., 2023) and decryptE dataset in Jurkat cells (Eckert et al., 2024) allowed us to compare consistent and inconsistent responses to HDACi treatment in non-isogeneic cellular backgrounds (Figure 1D). Proteome remodeling of HCT116 cells treated with vorinostat in our dataset were highly correlated with HCT116 cells treated with vorinostat in DeepCoverMOA (r_Pearson_ = 0.86, slope = 1.01, Figure 1E). Correlation between vorinostat-treated HCT116 cells in DeepCoverMOA and vorinostat treatments in our data was lower (r_Pearson_ = 0.50-0.69, Figure 1D).Consistent with DeepCoverMOA and decryptE, the abundance of CDKN1A, CCND3 and c-Jun increased with HDACi treatment (Richon et al., 1996; Vrana et al., 1999; Xiao et al., 1999), while the abundance of CCNA2 decreased with HDACi treatment (Richon et al., 1996; Xiao et al., 1999) (Figure 1E).

A549 cells harbor similar oncogenic KRAS and SMARCA4 mutations to HCT116 cells. When treated with vorinostat, A549 cells also had the highest correlation to HCT116 cells in the DeepCoverMOA vorinostat treatment (r_Pearson_ = 0.69). PC9 cells with vorinostat had the lowest correlation compared to the DeepCoverMOA data (r_Pearson_ = 0.50, Figure 1E). PC9 cells were also the most sensitive to HDACi perturbations (Figure 1B) and they did not have the increased abundance of JUN due to HDACi as seen in the other four cell lines (Figure 1E). Comparing our belinostat, TSA, and vorinostat treatment data and the decryptE analysis of HDACi treatment of Jurkat cells (Eckert et al., 2024), we observed lower overall correlation with the decryptE Jurkat cells responses to HDACi (r_Pearson_ = 0.15-0.50, Figure 1D). These data highlight the importance of considerations of the cellular context of cells when assessing mechanism of action and polypharmacology using proteomics.

**Figure 1.**
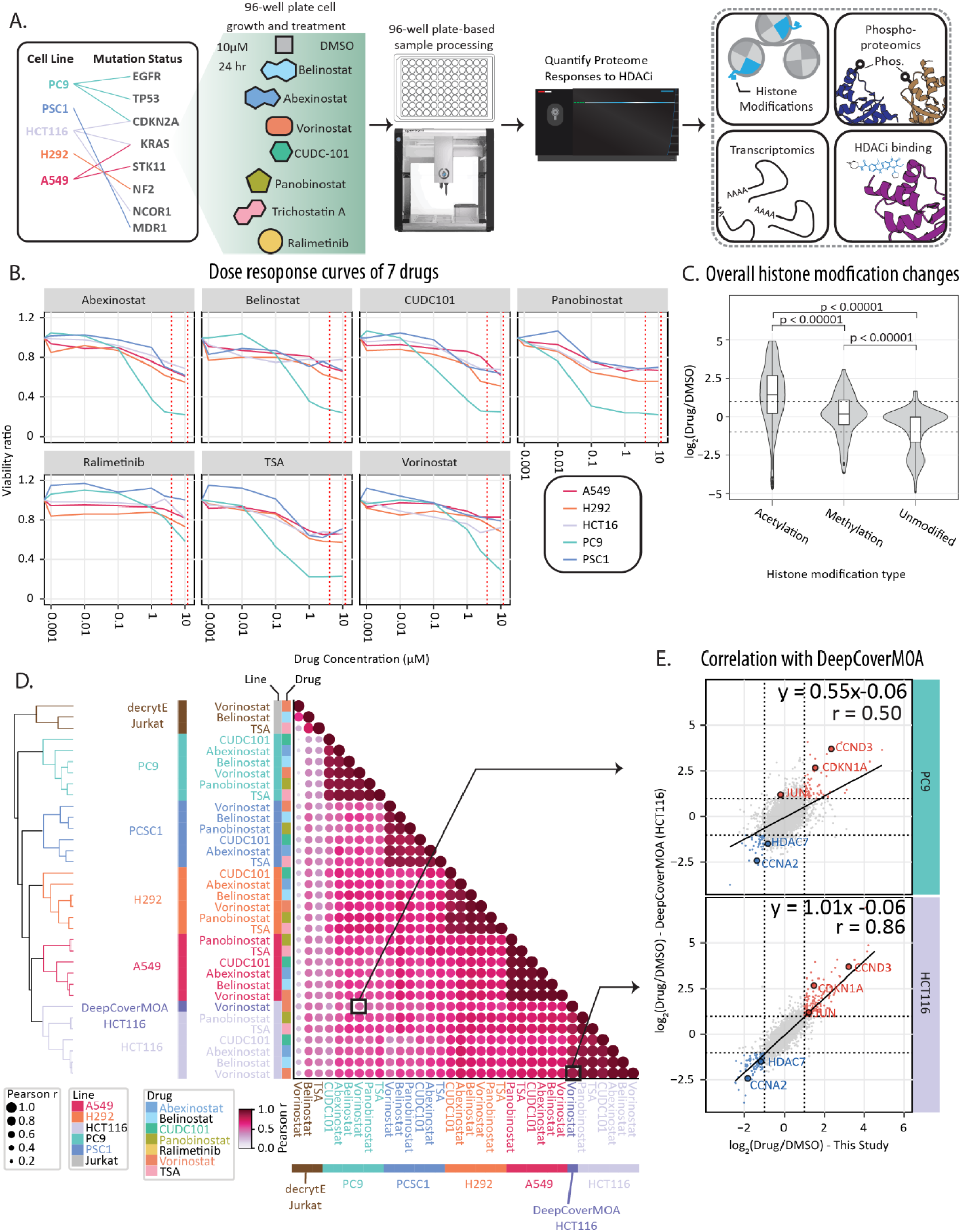
Heterogeneous responses in cancer cells. **A**, Overview of the workflow, starting with growing five cell lines in 96-well-plates. Sample preparation was performed with OT2 liquid handler after 24 h drug treatment. The measured Pearson correlation. **B,** Dose response curve of each drug tested on 5 cell lines normalized to DMSO treated controls. Viability was measured by luminescent cell viability assay. Viability ratio is drug treated cells/DMSO treated cells. **C,** Violin plot of histone modification abundance changes (center line: median; upper box limit: third quartile; lower box limit: first quartile; whiskers: 1.5x interquartile range; points: outliers). **D,** Pearson correlation analysis comparing HDACi treated samples in this dataset (10 μM HDACi, 24 hours, HCT116, A549, PSC1, PC9, H292), DeepCoverMOA (10 μM HDACi, 24 hours, HCT116), and decryptE (10 μM HDACi, 18 hours, Jurkat). **E**, Scatterplots highlight specific results from the correlation analysis for (top) the poorly correlated vorinostat-treated HCT116 cells from the DeepCoverMOA study compared to vorinostat-treated PC9 cells from this study and, (bottom) highly correlated vorinostat-treated HCT116 cells from the DeepCoverMOA study compared to vorinostat-treated HCT116 cells from this study. The log_2_FC is calculated by comparing drug treatment with DMSO.

### HDAC inhibition drives cell-type specific alteration of HDAC but not histone protein abundance landscape

In total, we observed 16,713 cell-line-by-drug regulated events across all groups on 2,553 different proteins (|log2FC| > 1, Supplementary Figure 2A, Supplementary data 1). Among the 2,553 regulated proteins, we quantified 2,282 proteins that were differentially abundant in less than 14 cell-line-by-drug groups—suggesting cell type- or cell state-specific responses (Supplementary Figure 2A). Based on the total number of regulated proteins, we observed the largest degree of proteome remodeling in HDACi-treated A549 cells (Figure 2A, Supplementary Figure 2B). Conversely, our outgroup treatment of cells with ralimetinib generated the lowest number of regulated proteins (Figure 2A). Chemoresistant PSC1 cells had the lowest number of regulated proteins among all tested cell lines which is consistent with the limited effect of drug treatment on cellular viability in these cells (Figure 2A, Supplementary Figure 2B). When comparing compound-specific effects, belinostat treatment of lung cancer cells (A549, H292, PC9 and PSC1) consistently generated the largest degree of proteome remodeling, followed by panobinostat, abexinostat, TSA, CUDC-101 and vorinostat (Figure 2A).

Across all five cell lines, we quantified 10 of the 11 human HDAC proteins in response to HDACi (Figure 2B). HDAC11 was not observed likely due to its generally low abundance in lung cells (Uhlén et al., 2015). We observed changes in both individual HDAC abundances and more generally in the responses of classes of HDACs. Class I HDACs include HDAC1, 2, 3 and 8, and these are generally found in the nucleus (Y. Li & Seto, 2016). Class II HDACs include HDAC4, 5, 6, 7, 9 and 10, which shuttle between the nucleus and cytoplasm (Y. Li & Seto, 2016). In our analysis, the abundance of Class I HDACs generally did not change after HDACi treatment, but we observed differential abundance for three out of six Class II HDACs (HDAC5/7/9, Figure 2B). HDAC7 protein abundance decreased across all cell lines, HDAC9 protein abundance decreased in PC9 and PSC1, while HDAC5 protein abundance increased in A549, PSC1 and HCT116, but decreased in PC9 (Figure 2B). Previous small-scale analyses found that HDAC7 (Class II), but not HDAC1 (Class I), protein abundance decreased with HDACi treatment (Caslini et al., 2019; Ma & D’Mello, 2011).

While HDAC protein abundances differed by class, the relative abundance of most histones did not have significant changes with HDACi treatment (Supplementary Figure 2C). Yet, we did find that histone modification status had cell line-specific HDACi-mediated responses (Figure 2C). Comparing A549 and H292 cells, the majority of histone modifications were consistent, e.g., the abundance of H3K14ac, H3K18ac and H3K4me1 increased with HDACi treatment and the abundance of H3K9me1, H3K9me2 and H3K9me3 decreased with HDACi treatment (Figure 2C). Conversely, H3K27ac abundance increased as expected in A549 cells treated with HDACi, but decreased in H292 cells. Similarly, H3K79me2 had increased abundance in H292 cells upon HDACi treatment, but reduced abundance in A549 cells (Figure 2C). Indeed, H3K79 methylation can antagonize modification of H3K27 (S. Chen et al., 2015). We note that the protein abundances of histone methyltransferases did not change in either cell line—e.g., for SETD2 which is associated with H3K36 methylation (Xie et al., 2022) or EZH1/EZH2 which are associated with H3K27 methylation (Nichol et al., 2016).

### Cell-type-specific regulation of histone modifications and chromatin regulatory factors with HDAC inhibition

In addition to HDAC and histone abundance, we investigated whether known histone binding proteins had differential protein abundance with HDACi treatment. Seven histone readers had abundance changes with HDACi treatment (Figure 2D), including histone chaperone ASF1B (histone H3 binder), histone reader BRD7 (Histone H3 binder), PHD finger proteins (PHF10/19/20/23), SWI/SNF chromatin remodeling protein ARID1B (Adhikary et al., 2016; Alvarez et al., 2011; Cong et al., 2006; N. Li et al., 2016; Mukherjee et al., 2020).

Among the 7 histone readers, ASF1B, BRD7 and PHF10 bind H3K14ac (Alvarez et al., 2011; Chugunov et al., 2021; Cong et al., 2006). The histone chaperone ASF1B recognizes H3K9me1 and imports H3 into the nucleus, where ASF1B can form a complex with H3K14K18ac (Alvarez et al., 2011). H3K9me1 abundance decreased in A549 cells treated with belinostat or vorinostat treatment, while H3K14ac and H3K18ac increased (Figure 2C, Supplementary Figure 2D). The abundance of PHF10 and BRD7 decreased with HDACi treatment, which was inverted from the phenotype we observed for ASF1B (Figure 2D). HDACi treatment resulted in increased H3K27ac in A549 cells, but decreased H3K27ac in H292 cells (Figure 2C, Supplementary Figure 2D). ASF1B overexpression is known to promote proliferation of cancer cells in a CDK9-dependent manner (X. Liu et al., 2020). Yet, we observed no significant change in CDK9 protein abundances (Supplementary Data 1). These data hint at a CDK9-independent route by which ASF1B stabilization in cancer cells may play a role in bypassing the effects of HDAC inhibition.

### Altered response of HDAC pathway members with HDACi treatment

Among the 2,553 proteins with significant HDACi-induced abundance changes, we found only 15 proteins with significantly altered abundances in all HDACi-treated cells (Figure 2E). We found this to be a striking result, but consistent with the poor correlation we observed with some previous datasets when comparing proteome responses after HDACi treatment (Figure 1E). The 15 proteins that changed consistently in all cell lines included known HDACi responsive proteins, such as increased protein abundance of the Wnt-signaling related protein DACT3 (Jiang et al., 2008) and the cell cycle regulatory protein cyclin-D3 (CCND3) (Richon et al., 2000; Siavoshian et al., 2000). More broadly, these 15 proteins were enriched for Wnt signaling and lung fibrosis annotations (Supplementary Figure 2E). We also found that several proteins previously annotated as HDACi responsive had significant abundance changes in a subset of our tested cell lines. These included increased abundance of cyclin-dependent kinase inhibitor 1 (CDKN1A) (Richon et al., 2000; Siavoshian et al., 2000); increased abundance of transcription factor and oncoprotein c-Jun (JUN)(Ma & D’Mello, 2011); decreased abundance of HDAC7 (Ma & D’Mello, 2011); and reduced abundance of cyclin-A2 (CCNA2) (Majumdar et al., 2012).

Based on these HDAC-class-specific changes, we sought to determine how HDACi treatment remodeled the canonical HDAC regulatory pathway components, such as the JUN signaling pathway. To explore the proteomic relationships surrounding JUN and HDACi treatments, we expanded the analysis to include direct interacting proteins of JUN based on the BioPlex interaction network (Huttlin et al., 2021; Schweppe et al., 2018) (Figure 2F). Within the JUN interaction network, the c-Jun transcription factor (JUN) had a significant increase in protein abundance in A549, H292, and HCT116 cells (log_2_FC_A549_ = 1.49; log_2_FC_H292_ = 1.43; log_2_FC_HCT116_ = 1.20), but not PC9 and PSC1 cells (log_2_FC_PC9_ = -0.09; log_2_FC_PSC1_ = -0.37; Figure 2F). We observed distinct differences in the abundance of the transcription factor JunB which decreased in H292 and PC9 cells (log_2_FC_H292_ = -0.84; log_2_FC_PC9_ = -1.1), but not in A549, HCT116 and PSC1 cells (log_2_FC_A549_ = 0.62; log_2_FC_HCT116_ = 0.30; log_2_FC_PSC1_ = 0.40, Figure 2F). We observed the same trend for c-FOS which forms a heterodimer with JUN and JunB, as c-FOS protein abundance significantly increased in A549, HCT116 and PSC1 cells (log_2_FC_A549_ = 1.82; log_2_FC_HCT116_ = 2.5; log_2_FC_PSC1_ = 3.62, Figure 2F).

We observed a negative correlation between JUN abundance and HDAC7 abundance and a positive correlation between JUNB and HDAC5 abundance (Figure 2G, H). However, there was no significant correlation between HDAC9 and JUN or JUNB. HDACi treatment induced decreased abundance of HDAC7 and increased abundance of JUN (Figure 2G). Since HDAC7 can inhibit JUN expression (Ma & D’Mello, 2011), these proteomic profiles suggest that decreased abundance, and thereby activity, of HDAC7 may be driving JUN abundance in response to HDACi. Moreover, the HDAC7 and JUN relationship was specific to HDACi-perturbed cells as no significant relationship between JUN and HDAC7 protein was observed in the large-scale CCLE proteomics screen of unperturbed cell lines (Nusinow et al., 2020), (Supplementary Figure 2F). HDACi treatment induced increased abundance of both HDAC5 and JUNB, a phenotype similar to that seen in which is similar to the injury-induced HDAC5 activity regulation of JUNB expression (Cho et al., 2013).

Beyond the relationship of JUN/JUNB with HDACi treatment, enrichment analysis (Liberzon et al., 2015) found that HDACi-responding proteins were significantly enriched for TNF-alpha signaling via NF-kB and UV response pathways (Figure 2I, J, Supplementary table 3). Additional pathways were only significantly regulated in a specific subset of cell lines. For example, mTORC1 associated proteins were only enriched in A549 and H292 cells treated with HDACi. Notably, this did not correlate with the background mutational status of these cell lines as A549 cells are driven by a constitutively-active KRAS mutant and H292 cells are driven by the loss of the NF2 tumor suppressor (van der Meer et al., 2019). Interestingly, proteins involved in p53 pathways mainly had increased abundance in A549 and HCT116 cells, while they had decreased abundance in H292 and PC9 cells, and these proteins were not differentially regulated in PSC1 cells.

**Figure 2.**
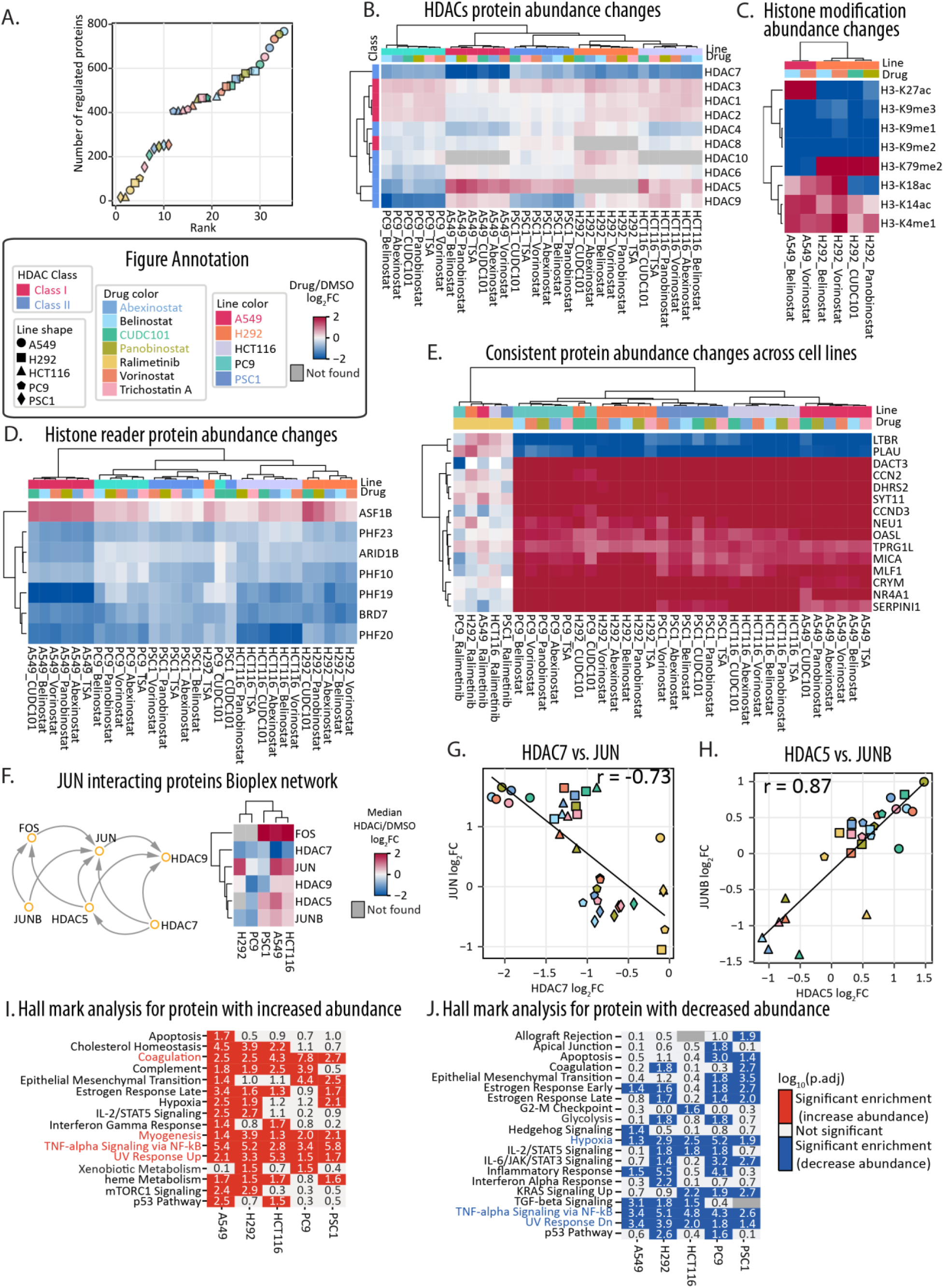
Altered response of HDAC and HDAC-related proteins. **A**, Waterfall plot highlighting the number of regulated proteins for each cell-line-by-drug group. **B,** Heatmap of protein abundance changes of 10 HDACs in 5 cell lines with HDACi treatment. **C,** Heatmap of histone modification abundance changes regulated in A549 and H292 with HDACi treatments. **D,** Heatmap of protein abundance changes of histone binding proteins in 5 cell lines with drug treatments. **E,** Heatmap of protein abundance changes of 15 proteins having abundance changes in all 5 cell lines with all HDACi treatments. **F,** Protein network from Bioplex between HDAC5/7/9 and JUN. The heatmap represents median protein abundance changes of HDAC5/7/9 and JUN with HDACi treatment in 5 cell lines. **G,** Comparison of protein abundance changes between HDAC7 and JUN. **H,** Comparison of protein abundance changes between HDAC5 and JUN. Pearson r is labeled on the top of the figure. **I, J,** Comparison of Hallmark enrichment analysis of regulated proteins from cell lines with HDACi treatment. Left with red color (**I**) is comparison of up-regulated proteins; right with blue color (**J**) is comparison of down-regulated proteins. Regulated proteins are proteins with absolute log_2_FC > 1. The p-value is calculated by comparing the observed frequency of an annotation term with the frequency that would be expected by chance. The p-value is adjusted by the Benjamini-Hochberg false discovery rate (FDR). Terms that have an adjusted p-value (p.adj) < 0.05 or log_10_(p.adj) > 1.3 are considered enriched. Both color scale and numbers represent log_10_(p.adj). For all figures above, the log_2_FC is calculated by comparing drug treatment with DMSO.

### Identifying direct effects of HDACi treatment on the proteome

After 24 hours of treatment with HDACi, the protein abundances for HDAC1 and HDAC2 did not change (Figure 2B), suggesting additional mediators of HDACi responses. To identify these mediators, we employed thermal proteome profiling to measure direct effects of HDACi treatment of A549 cells (Gaetani et al., 2019; Van Vranken, Li, Mintseris, Wei, et al., 2024) (Figure 3A). Owing to the potential importance of protein, cellular, and chromatin context in affecting HDACi engagement, we performed these assays in both cells and lysates to confirm protein-engagement data and determine likely direct versus indirect HDACi relationships (Van Vranken, Li, Mintseris, Wei, et al., 2024). In the presence of belinostat, HDAC1, HDAC2 and HDAC6, HDAC2 were significantly stabilized (Figure 3B).We also observed HDACi-mediated destabilization of HDAC1/2 containing complex components, including polycomb repressive complex components MGA and EHMT1/2 (Figure 3C, D) (Blackledge & Klose, 2021), and the BHC histone deacetylase complex component ZMYM3 (Hakimi et al., 2003; Leung et al., 2017).

In addition to thermal stability shifts for HDACs and HDAC-related proteins, HDACi altered the thermal stability of potential off-target proteins, including the cell cycle kinase AURKB (Figure 3B). AURKB was significantly destabilized with HDACi treatment in both cell- and lysate-based thermal stability analyses (Figure 3B). Re-analysis of bead-based small-molecule pulldown assays using HDAC-targeting pharmacophores (Lechner et al., 2022) revealed a dose-dependent response of AURKB to HDACi treatment similar to HDAC1/2/6, but not dose dependent response for the structurally similar AURKA (Figure 3E).

Measurement of A549 cellular responses confirmed that belinostat treatment increased AURKB-dependent phosphorylation of H3 Ser10 (Figure 3F, Supplementary Figure 3A). Addition of the AURKB inhibitor barasertib treatment, abolished H3 Ser10 phosphorylation. Consistently, phosphorylation on AURKB substrates and interacting partners INCENP and RACGAP1 increased with belinostat treatment (Supplementary Figure 3B). Based on these findings, we tested whether combination treatment of HDACi and AURKB inhibitors would further inhibit cell viability. Indeed, the combination of belinostat and barasertib treatment significantly reduced cell survival compared to either belinostat or barasertib alone (Figure 3G).

**Figure 3.**
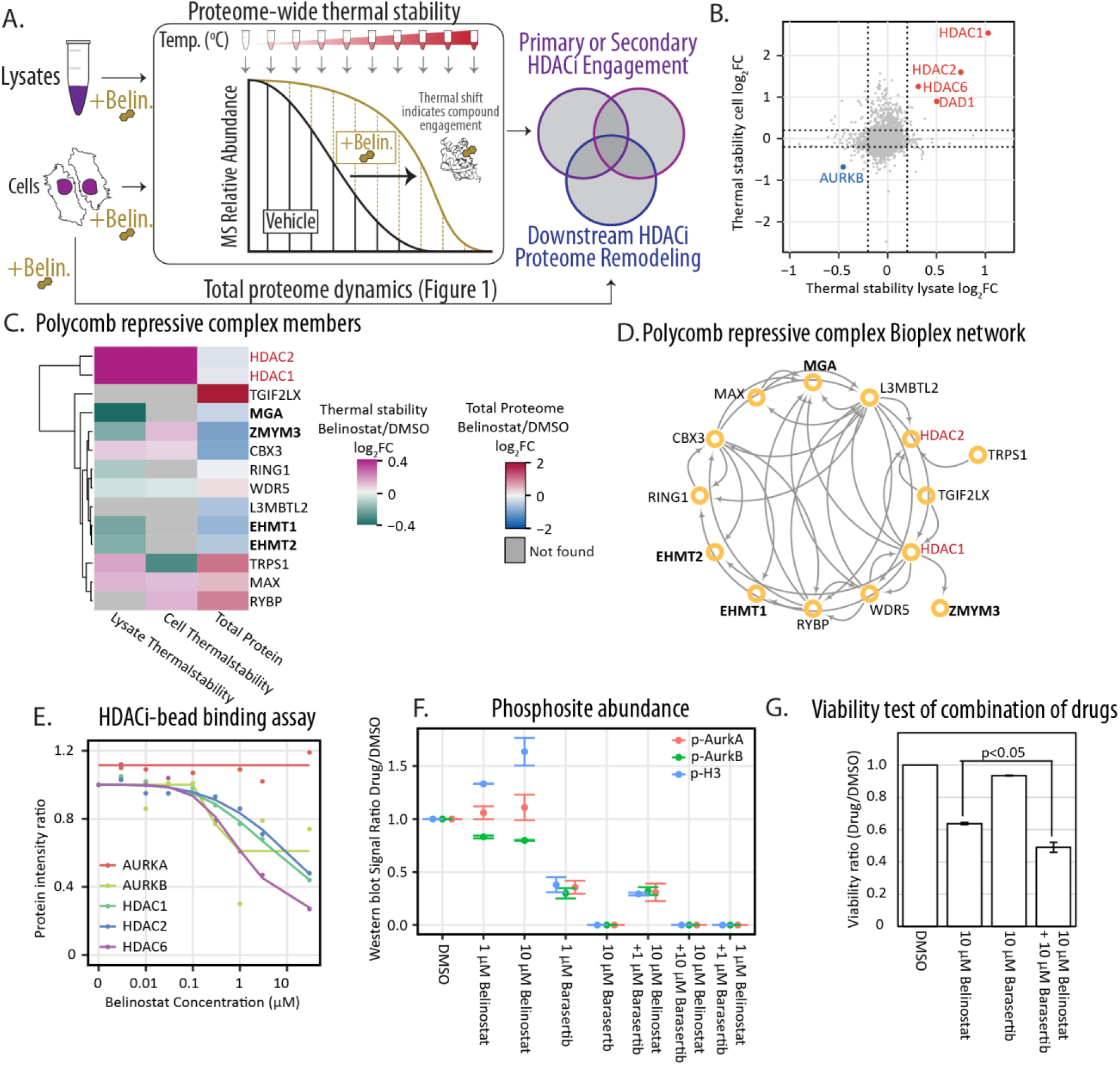
Analysis for protein-drug engagement for divergent cellular responses. **A**, Diagram shows cell and lysate thermal stability analyses and total proteome were performed using A549 cells or lysates treated with 10 μM belinostat. **B,** Comparison between two thermal stabilities of A549 with belinostat treatment. The log_2_FC is calculated by comparing drug treatment with DMSO. **C,** Heatmap of protein abundance changes in two thermal stability assay and total proteome responses of PRC1 members. **D,** Network from Bioplex of components of the polycomb repressive complex (PRC1). **E,** Dose-dependent response curve of AURKB to HDACi treatment from HDAC-targeting pharmacophores (Lechner et al., 2022). **F,** Western blot signal measured with Image J of AURKB, AURKA/B autophosphorylation and H3 ser10 phosphorylation with belinostat and barasertib treatment. Phospho-signals were normalized with the AURKB protein signal. Drug treatment signal was compared with DMSO treatment signal Western blot signal measurements **G,** Viability assay of A549 treated with combination of belinostat and barasertib.

### Co-regulatory analyses reveal a linkage between HDAC7 abundance and 14-3-3 protein interactions

In addition to direct effectors of heterogeneous HDACi response, we explored whether specific sets of proteins or protein complexes were co-regulated with HDACi treatment. Correlation analysis was performed comparing the 10 quantified HDACs and the 5,491 proteins quantified in all cell line-by-drug samples (Supplementary table 4). HDAC1 and HDAC2 are known to exhibit partially redundant effects in cells (Y. Zhang et al., 2023), and we observed that HDAC1 and HDAC2 protein abundances were correlated across datasets (r_Pearson_ = 0.92). Using this correlation approach, we observed 81 proteins with correlated protein abundances to at least one HDAC protein (r_Pearson_ > 0.9, Supplementary Figure 4A). In keeping with the cell line specific response of HDAC7 to HDACi treatment, we observed 83.9% of these proteins (68 of 81) were correlated with HDAC7 abundance (Supplementary Figure 4A). Of these proteins, the most highly correlated proteins (r_Pearson_ > 0.95) with HDAC7 were the tyrosine kinase SRC, MICALL1, and the fucosidase FUCA1 (Figure 4A, B, C).

The tyrosine kinase SRC is active in non-small cell lung cancers and involved in cellular transformation and metastasis (J. Zhang et al., 2007). Yet, consistent with our proteomics analysis, HDACi treatment can reduce SRC expression in cancer cells (Hirsch et al., 2006). While there was limited functional information about the role of MICALL1 (Molecule Interacting with CasL-like 1) in cancer, MICALL1 protein abundance in response to HDACi treatment was highly correlated with both HDAC7 (r_Pearson_ = 0.96) and SRC (r_Pearson_ = 0.90) protein abundance. In steady state protein abundance measurements across CCLE cancer cells, SRC, MICALL1, and FUCA1 had weak anticorrelation with HDAC7 (Supplementary Figure 4B, C, D). Thus, HDACi treatment seems to drive the coordinated protein abundances of these proteins potentially through shared interactions with 14-3-3 adapter proteins (YWHAH, YWHAG, YWHAB, YWHAZ, Figure 4D, 4E) (Golkowski et al., 2023; Göransson et al., 2006; Huttlin et al., 2021; Schweppe et al., 2018; Segal et al., 2023; Shen et al., 2020). The 14-3-3 proteins are known to regulate the nuclear import of Class IIa HDACs (HDAC4/5/7/9) and bind phosphorylated HDAC7 (X. Li et al., 2004; Nishino et al., 2008). Though many of the HDACi compounds used in our study primarily target both Class I and II HDACs (Y. Li & Seto, 2016), we found that the Class II HDAC7 exhibited robust, but cell-type-specific, responses to HDACi treatment across multiple lung cancer cells lines.

### HDACi alters phospho-signaling surrounding 14-3-3 regulated proteins

Based on the relationship between 14-3-3 proteins’ binding of phosphosites and their regulation of HDACi responses, we performed phosphoproteomics on HDACi treated cells. We identified a total of 6,611 phosphosites, including 2,271 phosphosites on 1,244 proteins with HDACi-dependent abundance changes (Supplementary data 2). Based on Kinase-Substrate Enrichment Analysis (KSEA) (Wiredja et al., 2017) of phosphosites with HDACi-depended abundance changes, activities for MARK2 and PRKAA2 were enriched in all cell lines. Activities for 8 additional kinases were enriched in all cell lines except PC9, and 9 kinase activities were enriched in at least one cell line (Figure 4F). PC9 cells treated with HDACi had the largest number of regulated sites, followed by H292, A549, HCT116 and PSC1 cells (Supplementary Figure 4E). PC9 cells were also the most susceptible to HDACi treatments in cell viability assays (Figure 1A). Cell lines treated with the p38 kinase inhibitor ralimetinib had the fewest regulated phosphosites (Supplementary Figure 4E).

The protein abundances of class II HDACs after HDACi treatment were highly correlated or anticorrelated with 27 phosphosites: HDAC5 (n_sites_=12), HDAC7 (n_sites_=11), and HDAC9 (n_sites_= 4) (r_Pearson_ > 0.8, quantified in at least 80% of our cell-line-by-drug groups, Supplementary Figure 4F). Related to the differential viability of cells in response to HDACi, the regulatory phosphosite Ser118 on the cell death regulator BAD was anticorrelated with HDAC7 abundance (r_Pearson_ = - 0.86, Figure 4G). Phosphorylation of BAD Ser118 disrupts Bcl-2/Bcl-XL binding inhibiting BAD-mediated apoptosis (J. Mann et al., 2019; Polzien et al., 2011). BAD Ser118 phosphorylation increased with HDACi in the three cell lines with intermediate sensitivity to HDACi (A549, H292, HCT116; Figures 1A, 4G). Conversely, no significant co-regulation was observed between HDAC7 and BAD protein abundance (r_Pearson_ = 0.13). BAD Ser99 phosphorylation, generally thought to act upstream of Ser118 phosphorylation, was also unchanged in response to belinostat (Supplementary data 2). Analysis of the decryptM phosphoproteomics responses confirmed vorinostat and abexinostat treatments increased BAD Ser118 phosphorylation with no change to Ser99 phosphorylation in MV4-11 and HeLa cells (Zecha et al., 2023). These data point to a cell-type-specific relationship between HDAC activity (and HDACi inhibition), HDAC abundance, and a BAD-dependent mechanism to overcome HDACi-mediated cell death.

Based on the consistent cell-type-specific responses, we sought to determine which kinases may be responsible for the HDACi-dependent phosphorylation of BAD. Previous work demonstrated that Ser118 can be phosphorylated by multiple kinases, including RAF, PKA, RSK, and AKT1 (Datta et al., 2000; Polzien et al., 2011) (Tan et al., 2000)(Datta et al., 2000). To determine which kinases may be responsible for Bad Ser118 phosphorylation, we performed coregulation analysis (Mitchell et al., 2023) between Ser118 abundance and the protein abundances of 248 protein kinases quantified in at least 28 HDACi-treated samples. In addition, based on the ‘RR-x-S’ motif surrounding BAD Ser118, we ran KinasePredictor (Poll et al., 2024). Among 77 kinases found in our data, AKT1 abundance changes had strong correlation with BAD Ser118 abundance changes (r_Pearson_ = 0.91), as well as a high KinasePredictor score (1.83). In addition to AKT1, there are three more kinases that have KinasePredictor score larger than 1 and protein abundance changes were correlated with BAD Ser118 abundance (absolute r_Pearson_ > 0.8). These included AKT2 (r_Pearson_ = - 0.87, Score_KinasePredictor_ = 1.93), ROCK1 (r_Pearson_ = -0.88, Score_KinasePredictor_ = 1.23) and SRPK2 (r_Pearson_ = 0.84, Score_KinasePredictor_ = 1.59) (Figure 4H).

**Figure 4.**
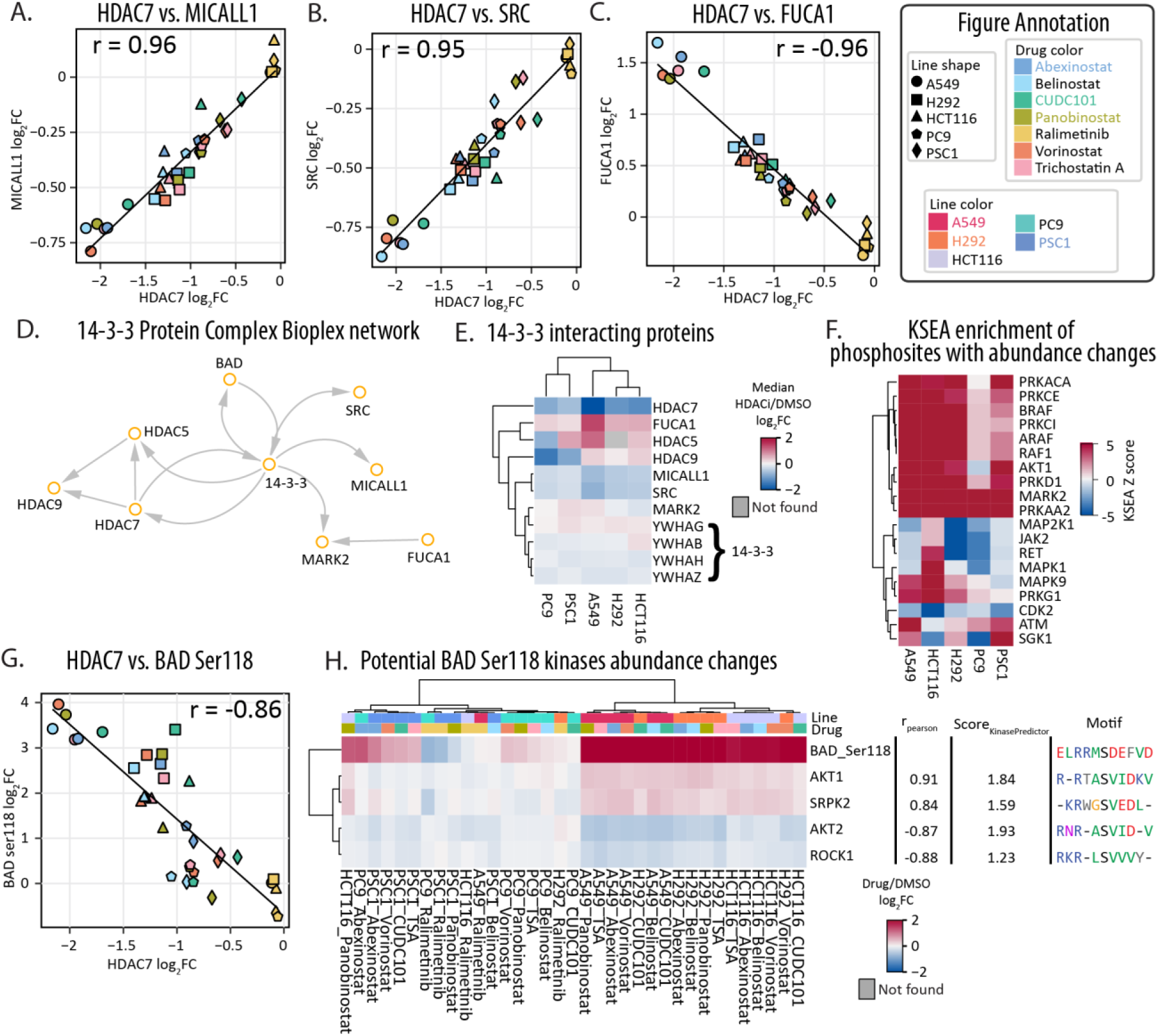
Altered response of 14-3-3 related proteins and phosphorylation. **A, B, C**, Comparison of protein abundance changes between HDAC7 and MICALL1 (**A**), SRC (**B**) or FUCA1 (**C**). Pearson r is labeled on the top of the figure. **D,** Network from Bioplex between 14-3-3 adapter proteins with HDAC5/7/9, MICALL1, SRC and BAD. **E,** Heatmap of median protein abundance changes of HDAC5/7/9 and 14-3-3 related proteins with HDACi treatment in 5 cell lines. **F,** KSEA enrichment of phosphosites having abundance changes with HDACi treatment in 5 cell lines. **G,** Comparison of abundance changes between HDAC7 and BAD ser118 phosphorylation. Pearson r is labeled on the top of the figure. **H,** Heatmap of phosphorylation abundance changes of BAD ser118 and protein abundance changes of correlated kinases. For all figures above, the log_2_FC is calculated by comparing drug treatment with DMSO.

### Coordinated remodeling of the transcriptomes and proteomes of HDACi perturbed cells

Owing to the inherent relationship between HDAC activity and transcription, we compared A549 drug-perturbed proteomics data to pseudo-bulked single-cell transcriptome with matched treatment conditions (abexinostat, panobinostat, belinostat, TSA) (Srivatsan et al., 2020), Supplementary Figure 5A). There were 232 transcripts with shared differential expression between protein and transcripts in all 4 treatments (Supplementary Figure 5B). In response to HDACi, A549 cellular proteomes and transcriptomes were generally correlated (r_Pearson_ = 0.48-0.64, Figure 5A), consistent with previous work (r_Pearson_ = 0.42-0.57) (Wang et al., 2019). We grouped protein and transcript responses to HDACi treatment into nine groups based on significant changes in proteomic and transcriptomic responses—e.g., Group 1 had increased transcript expression decreased protein abundance with HDACi treatment while Group 3 had concordant increased transcript expression and increased protein abundance with HDACi treatment (Supplementary Figure 5C).

Concordant transcript and protein responses to HDACi treatment in Groups 3 and 7 were observed for 505 protein/transcript pairs. As expected, the proteins in these groups were enriched for oncogenes and tumor suppressors, with notable effects of increased abundance of JUN, FOS and CDKN1A, and reduced abundance of HDAC7 and CCNA2 (Figure 5A). We then defined protein- or transcript-specific responders in Groups 1, 4, 6, and 9—i.e., those with differential abundance of proteins with HDACi treatment, and no effect on the transcripts, or vice versa (Figure 5A). With HDACi treatments, cyclin D3 (CCND3) was discordantly regulated with increased protein abundance after HDACi treatment and decreased transcript expression (Figure 5A). Conversely, the cyclin-dependent kinase inhibitor CDKN1A had a concordant relationship between its transcript and protein abundances with HDACi treatments (Figure 5A).

Since CCND3 coordinates cell cycle progression through binding and activation of CDK4 and CDK6 kinases, the discordant proteomics and transcriptomic profiles hint at a role of post-translational control of CCND3 protein abundance. One route for post-translational control of CCND3 could be through stabilization of the CDK4-CCND3 or CDK6-CCND3 protein complexes (LaBaer et al., 1997; Bockstaele et al., 2006; Sawai et al., 2012) (Figure 5B). We found that CDK6, but not CDK4, protein abundance decreased in A549 cells with HDACi treatment (Figure 5C). Consistent with DeepCoverMOA and decryptE data, we found that CDK6 protein abundance significantly decreased with HDACi treatment in the DeepCoverMOA and decryptE datasets (Figure 5D) (Eckert et al., 2024; Mitchell et al., 2023). However, CDK4 protein abundance was not significantly changed in our data or in DeepCoverMOA’s HCT116 study, but CDK4 abundance did decrease in Jurkat cells suggesting a route for heterogenous cell cycle responses to HDACi treatment (decryptE)(Figure 5D).

**Figure 5.**
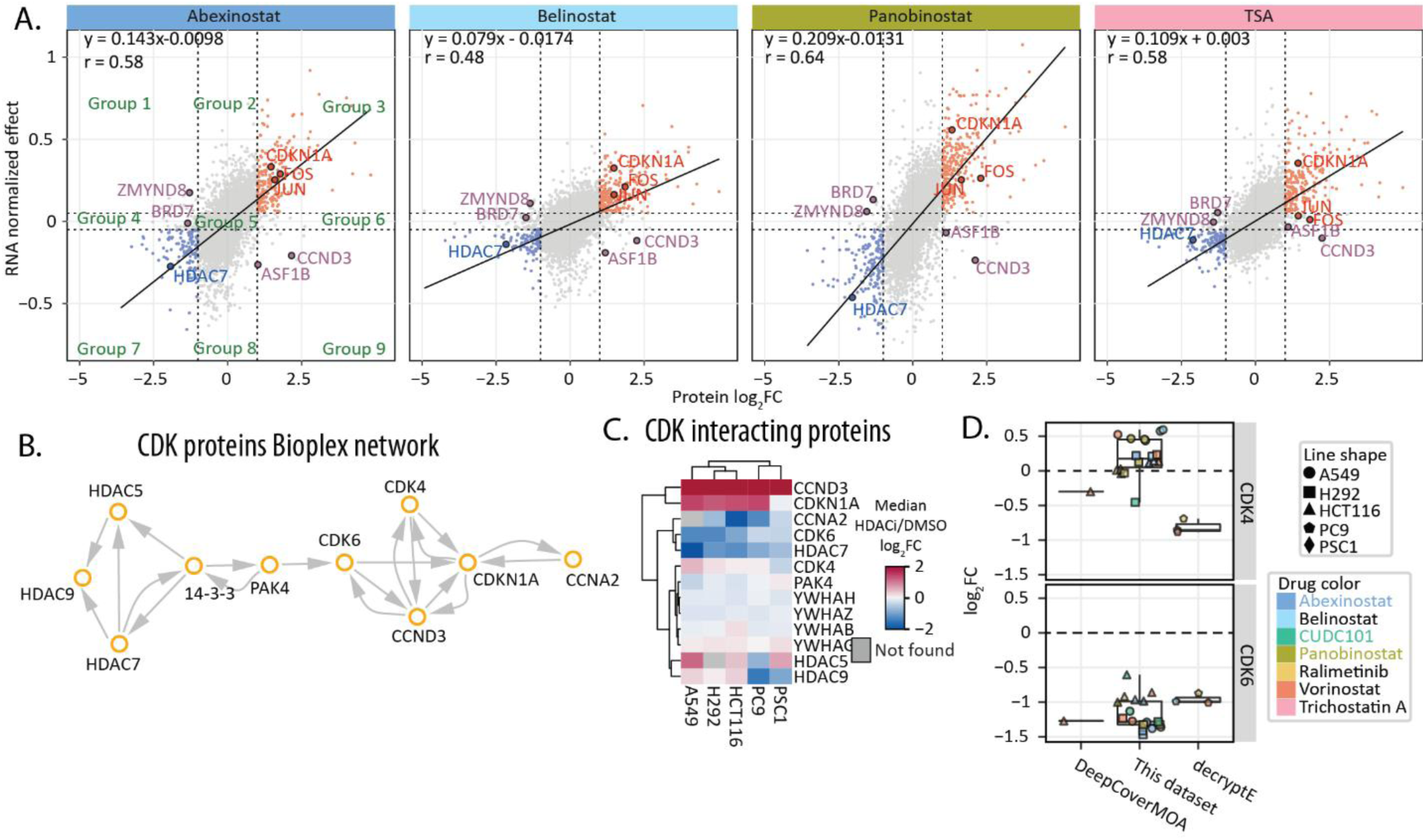
Coordination between transcriptomes and proteomics in A549 with HDACi treatments. **A**, Activity of A549 treated with 4 HDACi by the protein abundance log2FC (x-axis) compared with DMSO and RNA normalized effect (y-axis). The regulated proteins were defined as absolute log_2_FC > 1 in the proteomics dataset and absolute normalized effect > 0.05 in transcriptome dataset. Based on the four thresholds, proteins were grouped into 9 clusters. **B,** Network from Bioplex between 14-3-3 adapter proteins with HDAC5/7/9 and cyclin-dependent kinases. **C,** Heatmap of median protein abundance changes of 14-3-3 adapter proteins, HDAC5/7/9 and cyclin-dependent kinases with HDACi treatment in 5 cell lines. **D,** Boxplot of protein abundance changes of CDK4 and CDK6 in this dataset, DeepCoverMOA and decryptE with HDACi treatments. For all figures above, the log_2_FC is calculated by comparing drug treatment with DMSO.

## Discussion

Disparate cell states within cancer cells, driven by differential genetics, signaling axis activation, and protein interactions lie at the heart of inconsistent outcomes for therapeutic treatments. We investigated this phenomenon in the context of pleiotropic epigenetic (HDAC) inhibition as a model. HDAC inhibitors are actively used clinically for hematological malignancies and also promising preclinical activities for solid tumors, but with disputed efficacy based on cancer type and known adverse effects (Mamdani & Jalal, 2020). Our multiomic study reinforces that it is essential to characterize the molecular outcomes of therapeutic intervention that represent the diversity of molecular starting points (e.g., genetic backgrounds). In the context of lung cancer cell models, the majority of cell line models have a remarkable imbalance of race and gender, which call for action to establish omics resources to characterize these heterogeneous responses to drugs (Leon et al., 2023). Our work delineates the heterogeneity of proteome remodeling with drug treatment was consistent with heterogeneity in the cell line viability analysis and can offer important clues as to the differential susceptibility of these cell models to HDACi perturbation.

At the outset of this work, several large databases existed for the characterization of diverse human cancer cell lines, such as recent chemical genomic (Srivatsan et al., 2020) and proteogenomic characterization of CCLE cell lines (Ghandi et al., 2019; Nusinow et al., 2020). Here, we focused on measuring steady-state proteome remodeling of genomically diverse cancer cell lines after exposure to drug perturbation. We compared our lung cancer specific proteomics data with recent databases describing mechanistic annotation of drugs by profiling proteomics data, such as DeepCoverMOA (Mitchell et al., 2023), and the decrypt datasets (Eckert et al., 2024; Zecha et al., 2023). In shared cell-by-drug sample sets with these datasets, we observed consistent HDACi perturbation effects on the abundance of CDKN1A and CCND3 (Richon et al., 2000; Siavoshian et al., 2000); increased abundance of JUN protein and decreased abundance of HDAC7 (Ma & D’Mello, 2011); and reduced abundance of CCNA2 (Majumdar et al., 2012).

Our data from multiple cell lines also revealed new heterogeneity in HDACi treatment response. The protein abundance of JUN in A549, H292 and HCT116 cells, but not regulated in PC9 cells, correlated with PC9 cells’ poor viability with HDACi treatment. Upregulation of JUN is associated with cell proliferation and differentiation (Papavassiliou & Musti, 2020), which could explain PC9 is more sensitive to HDACi. Second, owing to the use of a single cancer cell line in many of these studies (Eckert et al., 2024; Mitchell et al., 2023), our data added a new level of understanding for how non-isogenic cell lines respond to chemical perturbation. Building on this work, the data presented here provides a molecular landscape for how cancer cell lines behave when they are treated with epigenetic inhibitors, and how this information can be leveraged to understand new vulnerabilities for therapeutic treatments.

We a web interface for the extensive amount of data to allow users to explore the data as a resource for understanding chemical perturbation of diverse cell types (https://github.com/SchweppeLab/HDAC-Perturbation-DataViewer). We explored the non-negligible differences in drug responses across various lung cancer cell lines through proteome profiling of five distinct lines by integrating proteomics, phosphoproteomics, and histone modification data. By combining proteomics with thermal stability data and transcript profiles, we identified specific protein and protein complex effectors of HDACi treatment in lung cancer cells (e.g., polycomb variant PCGF6). HDACi engagement data also revealed potential mechanisms of HDACi, such as through the engagement of DAD1 and AURKB. From our viability assay, A549 cells were more sensitive to belinostat than any other HDACi and this compound-specific sensitivity was not seen for the other KRAS mutant cell line (HCT116) or the other moderately sensitive H292 cells. Steady state AURKB protein abundance was 5.7-fold lower in A549 cells compared to H292 cells (Nusinow et al., 2020) potentially making A549 cells especially sensitive to off-target inhibition of AURKB.

Discordant changes of transcript and protein abundance revealed multiple potential routes by which cells can compensate for HDACi initiated cell death. For the histone reader ASF1B that can interact with H3K27ac and H3K79me2 (Jasencakova et al., 2010), we observed discordant transcript and protein abundance with HDACi treatment. Since the transcript abundance of ASF1B did not change significantly while protein abundance increased, we speculate that ASF1B protein abundance may be regulated post-translationally, potentially through hyperacetylation of H3 leading to nuclear sequestration and protection from degradation. In addition to the potential H3/ASF1B response to HDACi, our data show that abundance changes for specific histone modifications were correlated with protein and/or transcript abundance for cell cycle and chromatin regulators CCND3, ASF1B, PHF10 and BRD7. Since ASF1B, PHF10 and BRD7 can recognize H3K14ac. For BRD7 we note that only through the comparative transcriptomic and proteomic analysis did this protein become a strong candidate for future work owing to the moderate decrease in protein abundance but a discordant increased transcript abundance.

Finally, cell-line models are inherently reductionist and lack microenvironmental, immune, and pharmacokinetic contexts. Therefore, drug responses will not fully represent *in vivo* behavior. Given this limitation, by profiling multiple genetically diverse lung cancer lines using proteomics, phosphoproteomics, and histone mapping we were still able to reveal reproducible HDACi signatures. Together, the integration of our HDACi engagement data with previous data enabled identification of proteomic effectors of heterogenous cellular responses to HDACi. Thus, although this dataset focuses on HDAC inhibition, extension of our multiomics approach to include additional epigenetic drugs, larger cohorts of diverse cell lines, and improved methods to enhance throughput would further empower our understanding of key effectors of heterogeneous molecular responses in cell models.

## Methods

### Cell culture conditions

A549 (CVCL_0023), H292 (CVCL_0455), HCT116 (CVCL_0291), PC9 (CVCL_B260), and PSC1 (CVCL_5622) cells were dispensed into tissue culture treated 96 well plates at a density of 37,500 cells per well in a volume of 200 μl. After 24 h, cells growing at less than 80% confluence were then treated with 10 μM of each drug, including abexinostat, belinostat, CUDC-101, trichostatin A (TSA), panobinostat, vorinostat (SAHA), and ralimetinib, or 0.1% DMSO. All treatments were performed in 6 replicates: 2 replicates for proteome analysis, 2 replicates for viability analysis and 2 replicates for BCA assay. After treatment, cells were incubated for 24 h at 37 °C before washing with 200 μl PBS per well. After all PBS was aspirated, clear cell culture plates were immediately used for sample preparation for proteomics analysis or viability assay. Viability assay was performed with CellTiter-Glo® Luminescent Cell Viability Assay (Promega).

### Sample preparation for proteomics analysis

Sample lysis, reduction, alkylation, SP3 and digestion were processed with liquid handler OT2. Lysis buffer for BCA assay contains 8 M urea, 50 mM NaCl, 200 mM EPPS pH 8.5, Roche protease inhibitor tablets. Lysis buffer for proteomics analysis contains 5 mM TCEP in addition for protein reduction. Lysis buffer 30 μl was added into each well, followed by shaking at 500 rpm for 30 min. Protein concentrations were measured using Pierce BCA assay kits. Proteins for proteomics analysis were then alkylated with 30 μl 10 mM iodoacetamide for 30 min in the dark. The alkylation reaction was quenched by adding an additional aliquot of DTT. Proteins were then transferred to a 96-well PCR plate before being isolated using SP3. In brief, 2 μl of each bead type was added to each well before adding 65 μl neat ethanol and shaking for 30 min at 1000 rpm. Samples were placed on a magnet rack and supernatant was aspirated. Beads were washed with 125 μl 80% ethanol three times by resuspending. Beads were resuspended in 40 μl digest buffer (200 mM EPPS pH 8.5, 20 ng μl-1 LysC), then incubated overnight at 37 °C with constant agitation, before adding trypsin (400 ng) for an additional 6 hr at 37 °C. The final concentration of peptide was about 1 μg/μl. Two replicates of each cell line were manually labeled with a set of TMTpro 16-plex reagents, and the mixture was incubated at room temperature for 1 h. The reaction was quenched with 0.5% hydroxylamine, before pooling beads and eluates into a single 2 ml tube and placing it on a magnetic rack. Supernatant was collected. After labeling and mixing, peptide mixtures were desalted using C18 sep-pak cartridges (50 mg, Waters).

For phosphopeptide enrichment, PureCube magnetic beads were used with modified protocol (Leutert et al., 2019). In brief, peptides were dissolved in 80% ACN, 0.1% TFA and mixed with beads, shaking for 30 min. Phosphorylated peptides were eluted with 50% ACN, 2.5% NH4OH and then neutralized with 75% ACN, 10% formic acid. Elutes were desalted via Stage-tips prior to mass spectrometry analysis. Flow-throughs were then fractionated using basic-pH reverse phase chromatography (Navarrete-Perea et al., 2018). Briefly, peptides were resuspended in Buffer A (10mM ammonium bicarbonate, 5% acetonitrile, pH 8) and separated on a linear gradient from 13% to 42% Buffer B (10mM ammonium bicarbonate, 90% acetonitrile, pH 8) over an Agilent 300Extend C18 column using an Agilent 1260 HPLC equipped with wavelength detection at 214 nm and 254 nm). Fractionated peptides were desalted using Stage-tips prior to mass spectrometry analysis.

### Sample preparation for protein thermal stability assay

Thermal proteome profiling (TPP) by protein integral solubility assay (PISA) was performed as previously described (Van Vranken, Li, Mintseris, Gadzuk-Shea, et al., 2024). Each PISA experiment was performed in triplicate.

For live-cell PISA treatment, A549 cells were grown to a density of 1 x 10^6^ cells/mL, isolated by centrifugation, and resuspended in fresh media to 6 x 10^6^ cells/mL. to each well of a 24-well plate was added 500 uL of cell suspension and 1 mL of treatment (or vehicle) in media for a final cell concentration of 2x10^6^ cells/mL and a final treatment concentration of 10 µM. Cells were incubated with treatment for 1 hr at room temperature. 1 mL from each incubated sample (2x10^6^ cells) was collected, washed once in PBS, and resuspended in PBS containing 10 µM treatment. For each treatment condition, 30 µL of sample was aliquoted into 10 wells of a PCR plate and subjected to a temperature gradient from 48°C to 58°C for 3 minutes, then allowed to cool to room temperature for 5 minutes. 30 µL of ice-cold lysis buffer (200 mM EPPS pH 7.2, 150 mM NaCl, Roche cOmplete protease inhibitor, 0.5% NP-40) was added to each PCR plate well. 50 µL from each temperature point of a single condition was collected and pooled and cells were allowed to lyse at 4°C for 15 minutes. Lysate was spun at maximum speed for 2 hours and the soluble fraction was collected for sample preparation for proteomic analysis.

For lysate PISA treatment, cells were resuspended in the lysis buffer as above and lysed by repeated aspiration through a 21-gauge needle. Lysate was centrifuged at maximum speed for 30 minutes at 4°C. The soluble fraction was separated and lysate protein content was determined by BCA. Lysate and treatment were combined in lysis buffer for a final protein concentration of 1 mg/mL and a treatment concentration of 10 µM and then incubated at room temperature for 15 minutes. For each treatment condition, 30 µL of sample was aliquoted into 10 wells of a PCR plate and subjected to a temperature gradient as above. 25 µL from each temperature point of a single condition was collected, pooled and centrifuged at maximum speed for 2 hours. The soluble fraction was collected for sample preparation for proteomic analysis.

### Mass spectrometry data analysis for multiplexed quantitative proteomics

Peptides were separated prior to MS/MS analysis using an Easy-nLC (Thermo) equipped with an in-house pulled fused silica capillary column with integrated emitter packed with Accucore C18 media (Thermo). Mass spectrometric analysis was carried out on an Orbitrap Eclipse (Thermo). For whole proteome profiling, separation was carried out with 90-minute gradients from 98% Buffer A (5% ACN, 0.125% formic acid) to 28% Buffer B (90% ACN, 0.125% formic acid). Multiplexed analysis of samples was done using real-time search data acquisition (Schweppe et al., 2020), based on canonical SPS-MS3 acquisition. Briefly, a survey MS1 scan was used to identify potential peptide precursors (R = 120000, Mass range: 400-2000 m/z, max Inject time: 50ms, AGC: 200%, RF lens: 30%). The top 10 precursors were selected for fragmentation and analysis in the ion trap (Dynamic exclusion: 90s at 10 ppm, CID collision energy: 35%, max inject time: 50 ms, AGC: 200%, scan rate: rapid, isolation width: 0.5 m/z). Real-time spectral matching was carried out using the Comet search algorithm (Eng et al., 2013). If, and only if, a peptide was matched with high confidence, the instrument would then acquire an additional SPS-MS3 scan for quantification of relative abundances (R = 50000, HCD NCE: 45, max injection time: 150 ms).

For phosphopeptide profiling, separation was carried out with 90-minute gradients from 98% Buffer A (5% ACN, 0.125% formic acid) to 26% Buffer B (90% ACN, 0.125% formic acid). Multiplexed analysis of samples was done using canonical SPS-MS3 acquisition. Briefly, a survey MS1 scan was used to identify potential peptide precursors (R = 120000, Mass range: 300-2000 m/z, max Inject time: 50ms, AGC: 200%, RF lens: 30%). The top 10 precursors were selected for fragmentation and analysis in the ion trap (Dynamic exclusion: 90s at 10 ppm, HCD NCE: 30, max inject time: 35 ms, AGC: 250%, scan rate: rapid, isolation width: 0.5 m/z). SPS-MS3 was carried out in the Orbitrap (R = 50000, AGC: 250%, HCD NCE: 45, max injection time: 86 ms).

Raw spectral information was converted to mzXML format using Monocle (Rad et al., 2021), and spectra were matched using the Comet search algorithm compared against the Uniprot human database. TMTpro is a static modification at the N-terminus of peptides. The maximum missed cleavage was set as 2. Peptides and proteins were filtered to a 1% using rules of protein parsimony.

### Histone extraction and LC-MS/MS analysis for HDACi treatment

The histones were extracted and prepared for chemical derivatization and digestion as described previously (Bhanu et al., 2020; Sidoli et al., 2016). Briefly, lysine residues on histones were derivatized with the propionylation reagent (1:2 reagent: sample ratio) containing acetonitrile and propionic anhydride (3:1), with the solution pH adjusted to 8.0 using ammonium hydroxide. This propionylation reaction was performed twice and the samples were then dried using a speed vac. The derivatized histones were subsequently digested with trypsin at a 1:50 ratio (wt/wt) in 50 mM ammonium bicarbonate buffer at room temperature overnight. The N-termini of histone peptides were derivatized with the propionylation reagent twice and dried on speed vac (Searfoss et al., 2023). The peptides were desalted with the self-packed C18 stage tip. After desalting, the purified peptides were dried and reconstituted in 0.1% formic acid. Peptide analysis was performed using a LC-MS/MS system consisting of a Vanquish Neo UHPLC coupled to an Orbitrap Exploris 240 (Thermo Scientific). The histone peptide samples were kept at 7 °C on sample trays during LC analysis. Separation of peptides was carried out on an Easy-Spray™ PepMap™ Neo nano-column (2 µm, C18, 75 µm X 150 mm) at room temperature with a mobile phase. The chromatography conditions consisted of a linear gradient from 2 to 32% solvent B (0.1% formic acid in 100% acetonitrile) in solvent A (0.1% formic acid in water) over 48 minutes, followed by 42 to 98% solvent B over 12 minutes, at a flow rate of 300 nL/min. The mass spectrometer was programmed for data-independent acquisition (DIA) where each acquisition cycle consisted of a full MS scan, 35 DIA MS/MS scans of 24 m/z isolation width starting from 295 m/z to reach 1100 m/z. Typically, full MS scans were acquired in the Orbitrap mass analyzer across 290–1100 m/z at a resolution of 60,000 in positive profile mode with an auto maximum injection time and an AGC target of 300%. MS/MS data from HCD fragmentation was collected in the Orbitrap. These scans typically used an NCE of 30, an AGC target of 1000%, and a maximum injection time of 60 ms. Histone MS data were analyzed with EpiProfile 2.0 (Yuan et al., 2015) .

### Transcriptomics

Data regarding the HDACi-induced transcriptomic response in A549 was obtained from the NCBI GEO database under sample GSM4150378 (Srivatsan et al., 2020). Briefly, in this study, A549, MCF7, and K562 cells were treated with 4 increasing doses of a library of 188 small molecule compounds, including a subset of HDACi. The treated cells were subjected to high-throughput combinatorial indexing single-cell RNA-seq (sci-RNA-seq3). For this analysis, R package *monocle3* was used to analyze differentially expressed genes (DEGs) and for UMAP visualization (Trapnell et al., 2014; Qiu et al., 2017; J. Cao et al., 2019). For DEG analysis, the *celldataset* (cds) object was filtered to contain only A549 cells exposed to HDACi or vehicle control. For each HDACi, a generalized linear model was used to fit gene expression as a function of dose and replicate (quasi-Poisson regression) using *monocle3*’s *fit_models* function. This operation was performed on 7832 genes, determined as the union of genes expressed in 5% of cells per each inhibitor group. The p-values (Wald test) were FDR-corrected using the Benjamini-Hochberg method for multiple hypotheses, and the beta coefficients related to the contribution of dose were extracted and filtered as normalized beta coefficient > 0.05 and FDR < 5% for downstream comparison. Lastly, we performed batch integration (Haghverdi et al., 2018) and dimensional reduction (McInnes et al., 2018) of A549 cells treated with Abexinostat, Belinostat, Panobinostat, Trichostatin A, or vehicle to visualize cellular responses to HDACi.

### Data analysis

Protein quantification tables containing normalized TMT abundance ratios were analyzed with python. Coefficient of variation (CV) between replicates was calculated and filtered (CV < 0.3) before being calculated relative to DMSO. Networks were imported from BioPlex 3.0 (Schweppe et al., 2018; Huttlin et al., 2021) and modified in Cytoscape (Shannon et al., 2003). Molecular docking of belinostat (PubChem: 6918638) to AURKB (PDB: 4AF3 (Elkins et al., 2012)) was performed using AutoDock 4.2 within AutoDockTools 1.5.7 (Morris et al., 2009). Docking was performed using 100 Genetic Algorithm runs with Lamarckian output, all other parameters were set to default parameters.

Software used as listed:

| Software | Version |
| --- | --- |
| Python | 3.11.1 |
| pandas | 2.1.4 |
| numpy | 1.26.3 |
| matplotlib | 3.7.0 |
| plotnine | 0.12.4 |
| scipy | 1.11.3 |
| statsmodels | 0.14.0 |
| PyComplexHeatmap | 1.6.1 |
| seaborn | 0.13.0 |
| gsepy | 1.1.1 |
| scikit-learn | 1.3.1 |
| Cytoscape | 3.10.1 |
| BioPlex | 3.0 |
| AutoDockTools | 1.5.7 |
| AutoDock | 4.2 |

## Data availability

For total proteomics and phosphoproteomics, 70 .RAW files have been deposited to the ProteomeXchange Consortium via the PRIDE partner repository with the data set identifier PXD051206 (Username, password: DyNpQWlR). Spectra were searched against the human Uniprot database (download v.05/2020).

The data viewer can be found at https://github.com/SchweppeLab/HDAC-Perturbation-DataViewer. The data viewer was built in python using Streamlit and hosted through a docker container. Relevant information for starting the local server can be found in the GitHub repository README.

## Acknowledgement

We would like to acknowledge our funding sources: R35GM150919-01 (DKS), an Andy Hill CARE Distinguished Researcher Award (DKS), and a Cancer Consortium New Investigator Award (DKS). Support for this project was also provided by The Pew Charitable Trusts (DKS).

## Author contributions

C.L. and D.K.S. conceptualized the study. B.G., J.M. and D.K.S. supervised the experiments. C.L., C.S.,C.M.,R.K. and R.G. performed experiments. C.L. analyzed the data. C.L. drafted the original manuscript. Everyone reviewed and edited the manuscript. D.K.S. acquired funding.

