## Supplementary Figures for "Defining the heterogeneous molecular landscape of lung cancer cell responses to epigenetic inhibition"

**Supplemental data**

**Supplementary data 1** Normalized total protein abundance changes data

**Supplementary data 2** Normalized phosphosite abundance changes data

**Supplementary tables**

**Supplementary table 1** Table lists large-scale analysis of drug perturbations.

**Supplementary table 2** Information about the five cell lines used in this study

**Supplementary table 3** Hallmark, Go and KEGG pathway enrichment results of proteins with abundance changes

**Supplementary table 4** Pearson correlation table of HDACs with proteins that were quantified in all cell line-by-drug samples.

### Supplementary Figure 1

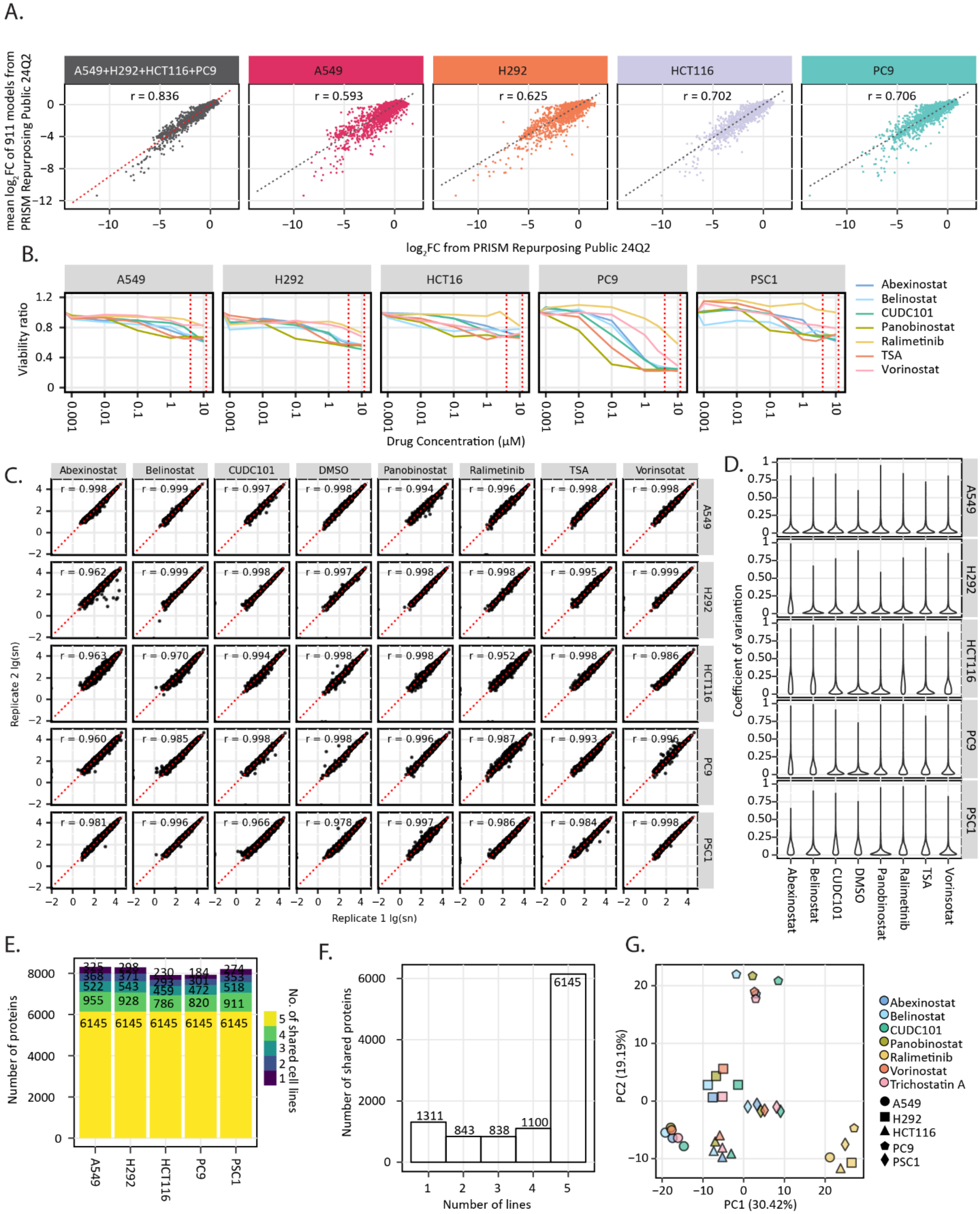


**Supplementary Figure 1 Heterogeneous responses in cancer cells and data reproducibility.** **A,** Comparison of viability of 4 cell lines, A549, H292, HCT116 and PC9, from PRISM drug repurposing public 24Q2. Each point represents one compound tested on the corresponding cell line. The x axis is log2 fold change of viability compared with negative control of each corresponding cell line. The y axis is the mean of log2 fold change of viability compared with negative control of 911 cell line models. **B,** Dose response curve of each cell line in response to 7 drugs normalized to DMSO treated controls. Viability was measured by luminescent cell viability assay. Viability ratio is drug treated cells/DMSO treated cells. For duplicate treatments of each cell line and drug combination, Pearson correlation (**C**) and coefficient of variations (**D**) of protein abundance changes were measured. The median Pearson r for these data was 0.996 and the median coefficient of variation was 4.4%. **E,** Total number of proteins found in each cell line. Different colors indicate the number of cell lines the protein was found in. Around 8000 proteins were found in each cell line, and more than 6000 proteins were shared by all 5 cell lines. **F,** Coverage of cell lines by proteins. 6145 proteins were found in all five cell lines, 1100 proteins were found in 4 cell lines, 838 were found in 3 cell lines, 843 were found in 2 cell lines and 1311 proteins were found in only one cell line. **G,** Principal component analysis (PCA) of 35 cell-line-by-drug groups. HDACi treated groups were clustered based on cell lines. All ralimetinib treated groups were clustered together (bottom right). Different shapes represent different cell lines. Different colors represent different drug treatments.

**Supplementary Figure 2**

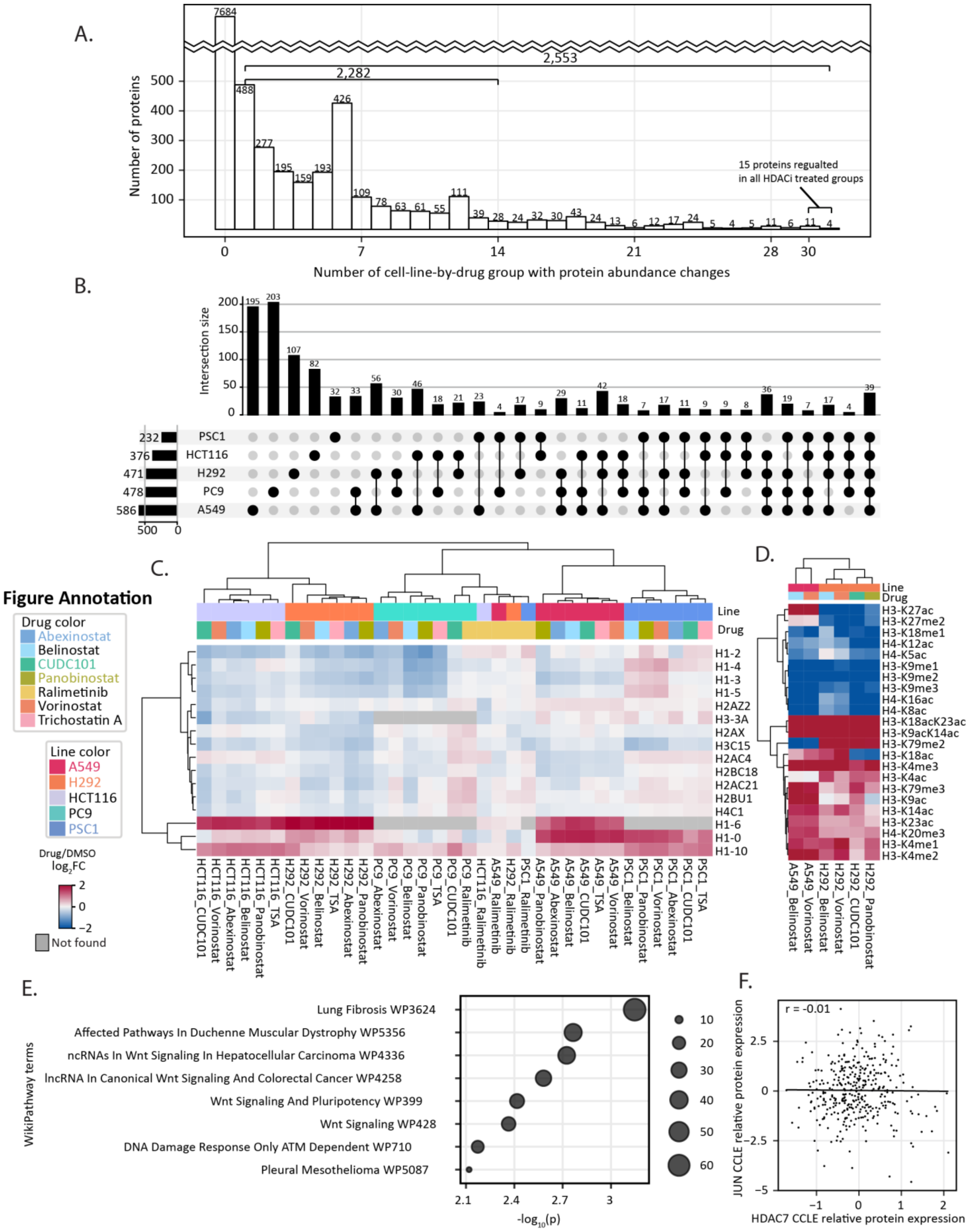


**Supplementary Figure 2 Altered response of HDAC and HDAC-related proteins. A,** Histogram of the number of regulated proteins shared by cell-line-by-drug groups. Regulated proteins are proteins with absolute $log_2FC > 1$. 7,684 proteins were not regulated in any cell-line-by-drug group. None protein was regulated in all 35 cell-line-by-drug groups, but 21 proteins were regulated in more than 28 cell-line-by-drug groups, 99 proteins were regulated in more than 21 cell-line-by-drug groups. **B,** Overlap between five lines of regulated proteins. 39 proteins were found regulated in all cell lines. 195 proteins were only regulated in A549, 203 only in PC9, 107 only in H292, 82 only in HCT116 and 32 only in PSC1. Regulated proteins are proteins with absolute $log_2FC > 1$. Proteins that are regulated with at least one drug treatment are defined as regulated in this cell line. **C,** Heatmap of protein abundance changes of histone proteins in 5 cell lines with drug treatments. **D,** Heatmap of histone modification abundance changes regulated in A549 and H292 with HDACi treatments. **E,** WikiPathway enrichment of 15 proteins having abundance changes in all 5 cell lines with all HDACi treatments. **F,** Pearson correlation of CCLE relative protein expression between HDAC7 and JUN. For all figures above, the $log_2FC$ is calculated by comparing drug treatment with DMSO.

### Supplementary Figure 3

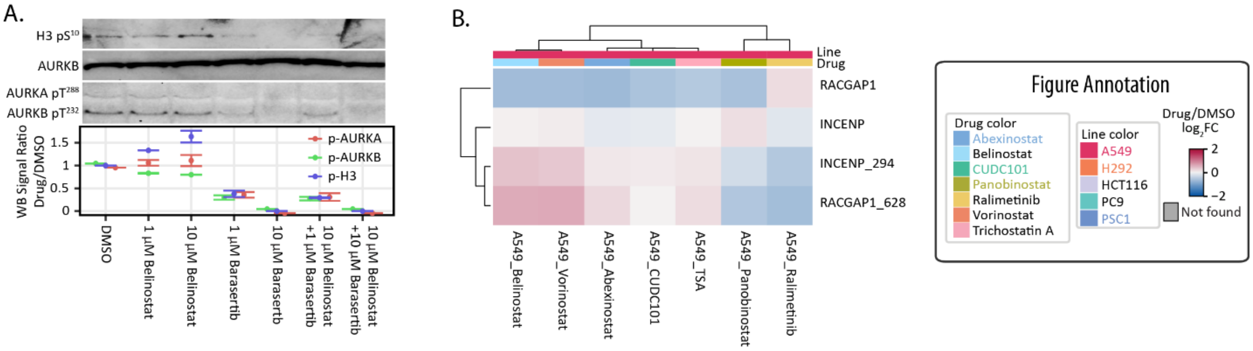


**Supplementary Figure 3 Belinostat affects AURKB activity. A,** Western blot shows AURKB, AURKA/B autophosphorylation and H3 ser10 phosphorylation with belinostat and barasertib treatment. Western blot signal measured with Image J. **B,** Heatmap of phosphorylation abundance changes and protein abundance changes of two AURKB interacting proteins, INCENP and RACGAP1.The log2FC is calculated by comparing drug treatment with DMSO.

### Supplementary Figure 4

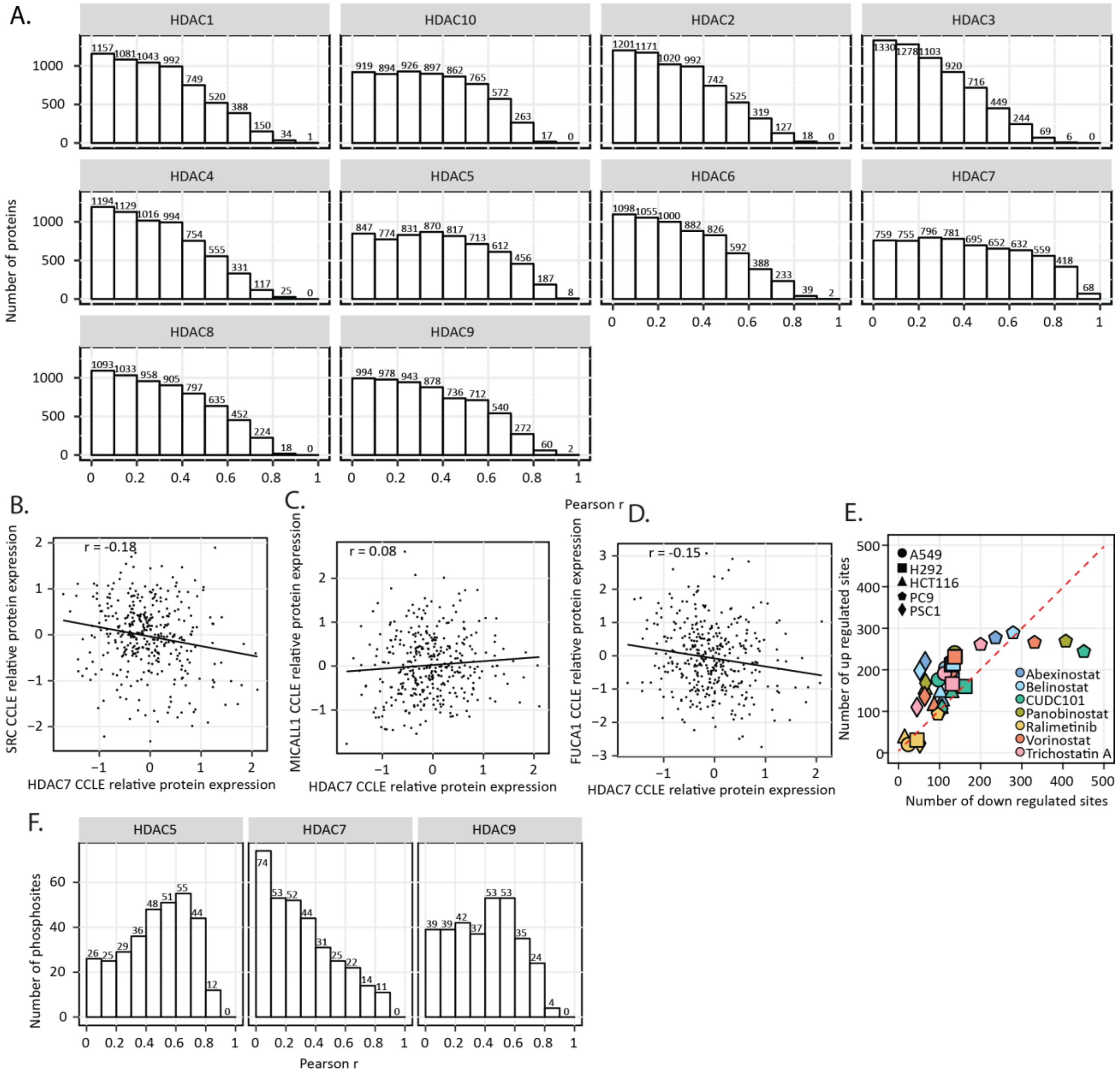


**Supplementary Figure 4 Altered response of 14-3-3 related proteins and phosphorylation. A,** Pearson correlation between HDACs protein abundance and other protein abundance. The x axis is the pearson r and y axis is the number of proteins. **B, C, D,** Pearson correlation of CCLE relative protein expression between HDAC7 and SRC (**B**), MICALL1 (**C**) or FUCA1 (**D**). **E,** Activity of each cell-line-by-drug group measured by the number of phosphosites down-regulated (x axis) and up-regulated (y axis) after 24 h treatment. Regulated phosphosites are phosphosites with absolute $log_2FC > 1$. **F,** Number of phosphosite abundance changes correlated with HDAC5, HDAC7 and HDAC9 protein abundance changes. For all figures above, the $log_2FC$ is calculated by comparing drug treatment with DMSO.

### Supplementary Figure 5

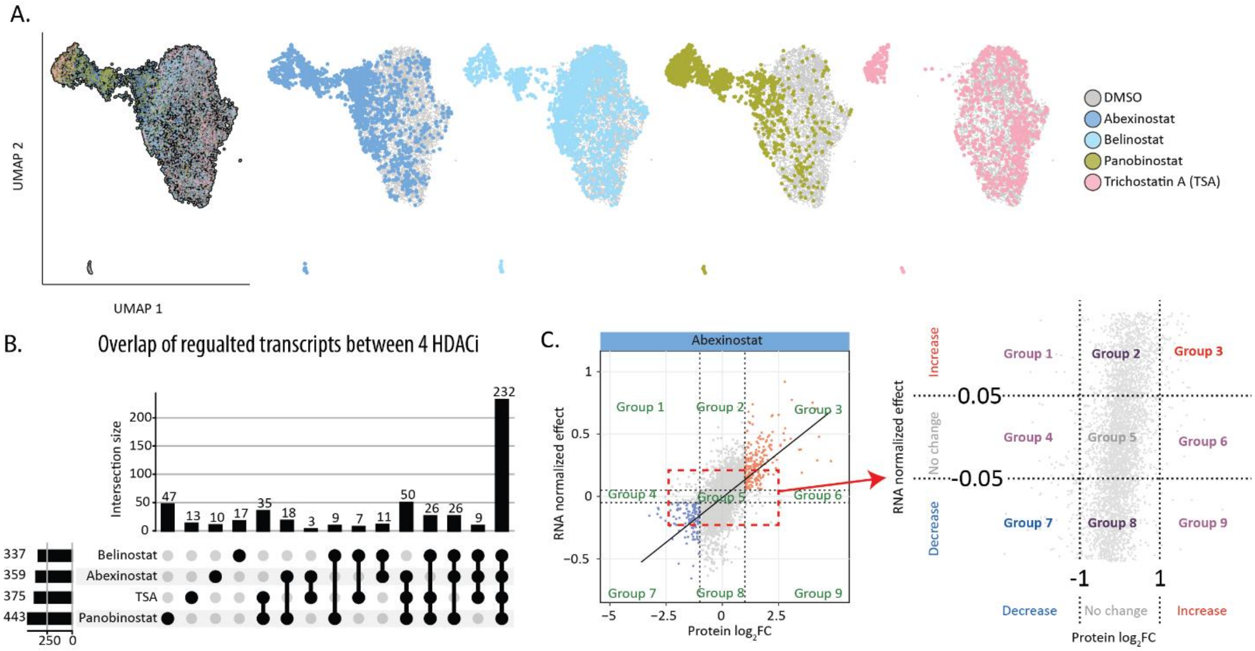


**Supplementary Figure 5 Coordination between transcriptomes and proteomics in A549 with HDACi treatments. A,** UMAP analysis of single-cell transcriptomics in A549 with 4 HDACi, abexinostat, belinostat, panobinostat and TSA. **B,** Upset plot shows number of overlap regulated transcripts shared between 4 HDACi. **C,** Diagram shows 9 groups based on transcript and protein changes. Proteins are divided into decrease ($log_2FC < -1$), no change ($-1 <= log_2FC <= 1$) and increase ($log_2FC > 1$) groups. The log2FC is protein abundance change compared with DMSO. Transcripts are divided into decrease (normalized effect < -0.05), no change (-0.05 <= normalized effect <= 0.05), increase (normalized effect > 0.05). The normalized effect is RNA relative abundance compared with vehicle control.
